# Functional coupling between NMDA receptors and SK channels in rat hypothalamic magnocellular neurons: altered mechanisms during heart failure

**DOI:** 10.1101/759720

**Authors:** HC Ferreira-Neto, JE Stern

## Abstract

Glutamatergic NMDA receptors (NMDAR) and small conductance Ca^2+^-activated K^+^ channels (SK) are critical synaptic and intrinsic mechanisms that regulate the activity of hypothalamic magnocellular neurosecretory neurons (MNNs) under physiological and pathological states, including lactation and heart failure (HF). Still, whether NMDARs and SK channels in MNNs are functionally coupled, and whether changes in this coupling contribute to exacerbated neuronal activity during HF is at present unknown. In the present study, we addressed these questions using patch-clamp electrophysiology and confocal Ca^2+^ imaging in a rat model of ischaemic HF. We found that in MNNs of sham rats, blockade of SK channels with apamin (200 nM) significantly increased the magnitude of an NMDAR-evoked current (I_NMDA_). We also observed that blockade of SK channels potentiated NMDAR-evoked firing, and abolished spike frequency adaptation in MNNs from sham, but not HF rats. Importantly, a larger I_NMDA_-ΔCa^2+^response was observed under basal conditions in HF compared to sham rats. Finally, we found that dialyzing recorded cells with the Ca^2+^ chelator BAPTA (10 mM) increased the magnitude of I_NMDA_ in MNNs from both sham and HF rats, and occluded the effects of apamin in the former. Together, our studies demonstrate that in MNNs, NMDARs and SK channels are functionally coupled, forming a local negative feedback loop that restrains the excitatory effect evoked by NMDAR activation. Moreover, our studies also support a blunted NMDAR-SK channel coupling in MNNs of HF rats, standing thus as a pathophysiological mechanism contributing to exacerbated hypothalamic neuronal activity during this prevalent neurogenic cardiovascular disease.

## INTRODUCTION

The neurohypophysial hormones vasopressin (VP) and oxytocin (OT) play a pivotal role in the regulation of several physiological processes such as regulation of vascular tone, baroreflex modulation, water reabsorption, sodium balance, reproduction, parturition and lactation (Poulain *et al*., 1977; Antunes-Rodrigues *et al*., 2004). VP and OT are synthesized and released by the magnocellular neurosecretory cells (MNNs) located in the supraoptic (SON) and paraventricular nuclei (PVN) of the hypothalamus (Swanson & Sawchenko, 1983).

Besides its important involvement in these physiological processes, VP has been shown to contribute to development and maintenance of prevalent cardiovascular diseases such as hypertension and congestive heart failure (HF) (Chatterjee, 2005; Kim *et al*., 2013; Ribeiro *et al*., 2015). Several studies have reported that human patients and animal models with HF exhibit chronically elevated plasmatic VP levels (Goldsmith *et al*., 1983; Riegger *et al*., 1985; Francis *et al*., 1990), which contribute to arterial vasoconstriction, fluid imbalance, persistent hypernatremia and kidney damage, all of which contribute to morbidity and mortality in HF patients (Cohn *et al*., 1981; Goldsmith *et al*., 1986a; Goldsmith *et al*., 1986b; Packer *et al*., 1987; Hodsman *et al*., 1988; Rouleau *et al*., 1994; Schrier & Abraham, 1999; Nakamura *et al*., 2000). Despite the growing evidence supporting neurohumoral hyperactivation and increased plasmatic VP concentration in HF, there is still little information regarding the precise mechanisms involved in the regulation of MNNs during HF.

The release of VP and OT into the bloodstream is directly regulated by the degree and pattern of the firing activity of MNNs, which is turn controlled by the combination of both intrinsic membrane properties, local modulators and synaptic inputs (Bourque *et al*., 1993; Bourque, 2008). Among the latter, glutamate is the main excitatory neurotransmitter in the hypothalamus, acting most predominantly on ionotropic N-methyl-D-aspartate (NMDA) receptors (NMDARs) (Hu & Bourque, 1992; Nissen *et al*., 1995; Li *et al*., 2003). In fact, numerous studies support a contribution of an altered glutamate signaling to neurohumoral activation in HF, including changes in the degree of afferent synaptic activity (Han *et al*., 2010; Potapenko *et al*., 2011), changes in the expression of postsynaptic receptors, including NMDARs (Li *et al*., 2003), and enhanced extrasynaptic NMDAR activation due to an impaired astrocytic glutamate uptake (Potapenko *et al*., 2012, 2013; Stern & Potapenko, 2013). Still, a comprehensive understanding of how these changes contribute to altered neuronal function and consequently neurohumoral activation during HF is still missing.

Gating of NMDARs results in an influx of Ca^2+^, which not only evokes a direct membrane depolarization, but also increases intracellular Ca^2+^ levels ([Ca^2+^]_i_) leading to activation of intracellular cascades and/or Ca^2+^-dependent ion channels (McBain & Mayer, 1994; Petersen, 2002; Sah & Faber, 2002), including the small conductance Ca^2+^-activated K^+^ channel (SK channels). Recent studies in hippocampal and cortical neurons reported that NMDAR activation leads to activation of different K^+^ channels, including SK channels to regulate synaptic transmission and plasticity (Ngo-Anh *et al*., 2005; Babiec *et al*., 2017).

SK channels in MNNs mediate an after-hyperpolarizing potential (AHP) that underlies spike frequency adaptation, thus strongly influencing repetitive firing properties in MNNs (Stern & Armstrong, 1997; Teruyama & Armstrong, 2002; Greffrath *et al*., 2004). Importantly, we recently demonstrated a diminished expression and function of SK channels in MNNs from HF rats (Ferreira-Neto *et al*., 2017), and changes in SK channel function/expression has been previously reported in other prevalent cardiovascular-related diseases, including salt-sensitive hypertension (Chen *et al*., 2010; Pachuau *et al*., 2014) and chronic high salt intake (Chapp *et al*., 2017). Still, whether a functional coupling between NMDARs and SK channels influences membrane excitability and firing activity in MNNs, and whether a change in this coupling contributes to altered neuronal activity during HF remains to be determined.

## MATERIALS AND METHODS

### Ethical Approval

Male Wistar rats (180-200 g), purchased from Envigo Laboratories (Indianapolis, IN), and male homozygous transgenic eGFP-VP Wistar rats (180-200 g) (Ueta *et al*., 2005) were used in this study. Rats were housed at room temperature (24–26°C) in a room with a 12:12-h light-dark cycle with normal rat chow and drinking water *ad libitum.* All procedures were approved by the Georgia State University Institutional Animal Care and Use Committee (IACUC) and carried out in agreement with the IACUC guidelines.

### Animals and induction of HF

HF was induced by coronary artery ligation as previously described (Biancardi *et al*., 2011). Briefly, animals were anesthetized with isoflurane (2-4%) and intubated for mechanical ventilation. A left thoracotomy was performed and the heart exteriorized. The ligation was placed on the main diagonal branch of the left anterior descending coronary artery. Buprenorphine SR-LAB (0.5 mg/kg sc; Zoo Pharm, Windsor, CO, USA) was given immediately after surgery to minimize postsurgical pain. Sham animals underwent the same procedure, but the coronary artery was not occluded. Transthoracic echocardiography (Vevo 3100 systems; Visual Sonics, Toronto, ON, Canada) was performed 4 wk after surgery under light isoflurane (3-4 %) anesthesia. All animals were used 6–8 wk after surgery. The left ventricle internal diameter, as well as the left diameter of the ventricle posterior and anterior walls, was obtained throughout the cardiac cycle from the short-axis motion imaging mode. Measured parameters were used to calculate ejection fraction and fractional shortening.

### Hypothalamic slice preparation

Hypothalamic brain slices were prepared according to methods previously described (Ferreira-Neto *et al*., 2015). Briefly, rats were deeply anesthetized with pentobarbital (80 mg/kg, i.p.). Then, rats were quickly decapitated; brains dissected out and coronal slices cut (240 μm thick) using a vibroslicer. An oxygenated ice-cold artificial cerebrospinal fluid (aCSF) was used during slicing (containing in mM: 119 NaCl, 2.5 KCl, 1 MgSO_4_, 26 NaHCO_3_, 1.25 NaH_2_PO_4_, 20 d-glucose, 0.4 ascorbic acid, 2 CaCl_2_, and 2 pyruvic acid; pH 7.4; 295 mOsm). Slices were placed in a holding chamber containing aCSF and kept at room temperature until used.

### Patch-clamp electrophysiology

Slices were bathed with an aCSF (~2.0 mL/min) containing 5 mM CsCl to suppress the depolarizing action potential and better isolate the AHP (Ghamari-Langroudi & Bourque, 1998; Teruyama & Armstrong, 2005) and were continuously bubbled with 95% O_2_–5% CO_2_ and maintained at ~30-32°C. Thin-walled (1.5 mm od, 1.17 mm id) borosilicate glass (G150TF-3, Warner Instruments, Sarasota, FL, USA) was used to pull patch pipettes (3–5 MΩ) on a horizontal Flaming/Brown micropipette puller (P-97, Sutter Instruments, Novato, CA, USA). The internal solution contained in mM: 135 KMeSO_4_, 5 EGTA, 10 HEPES, 10 KCl, 0.9 MgCl_2_, 4 MgATP, 0.3 NaGTP, and 20 phosphocreatine (Na^+^); pH was adjusted to 7.2–7.3 with KOH. The osmolality of the intracellular solutions was 285 mOsm. Recordings were obtained with an Axopatch 700A amplifier (Axon Instruments, Foster City, CA, USA) from magnocellular neurosecretory neurons (MNNs) from the SON using an infrared differential interference contrast (IR-DIC) videomicroscopy. The voltage output was digitized at 16-bit resolution, 10 kHz and was filtered at 2 kHz (Digidata 1440A, Axon Instruments). Data were discarded if the series resistance was not stable throughout the entire recording (>20% change).

Voltage-clamp recordings were performed to study the NMDA-evoked current (I_NMDA_). A puff of NMDA of varying durations (50 μM, 100-500 ms) was focally delivered to the patched neurons that were held at −50 mV. In order to block SK channels, the selective blocker apamin (200 nM) was bath-applied for 10 min. In some experiments, a subset of MNNs were dialyzed with the Ca^2+^ chelator BAPTA (10 mM) added to the recording pipette.

For current clamp experiments, MNNs were held at −50 mV and subjected to either direct current injection or a puff of NMDA, in order to evoke spiking activity. To evaluate the magnitude of spike frequency adaptation during the evoked trained of action potentials, we measured the interspike interval (ISI) for each evoked action potential in the train and normalized them to the first ISI of the train. Plots of normalized ISI as a function of the spike number in the train were generated in response to direct current injection or a puff of NMDA, before and after apamin application. The slope of a linear regression fitted to the plot (mean r^2^: sham 0.72 ± 0.06; HF 0.72 ± 0.07) was used to quantify the degree of SFA in each group and condition. For this analysis, and to be consistent and able to compare SFA between direct current injection and NMDAR-evoked firing, we focused on the first 10-15 spikes of the evoked train, which represented approximately the number of action potentials evoked during the puff of NMDA, excluding then from the analysis those spikes that occurred after the stimulation stopped, whose decrease in frequency represent the progressive washout of the drug. In a subset of experiments, the Na^+^ channel blocker tetrodotoxin (TTX, 1 μM) was bath-applied as indicated.

### Confocal calcium imaging

Neurons were loaded through the patch pipette with Fluo-5F pentapotassium salt (100 μM; Invitrogen) added to the recording solution, as previously described (Sonner *et al*., 2011). Imaging was conducted using the Andor Technology Revolution system (iXON EMCCD camera with the Yokogawa CSU 10, confocal scanning unit). Fluorescence images were acquired at a rate of 4 Hz, using an excitation light of 488 nm and emitted light at >495 nm. The fractional fluorescence (F/F0) was determined by dividing the fluorescence intensity (F) within a region of interest drawn within the MNN cell body after the NMDA focal application (50 μM, 500 ms) by a baseline fluorescence value (F0) determined from 50 images before the NMDA puff. Data were analyzed using ImageJ software.

### Statistical analysis

All values are expressed as mean ± standard error mean (SEM). Paired or unpaired Student’s *t* test and one- or two-way analysis of variance (ANOVA) tests with Bonferroni *post hoc* tests were used as indicated. Differences were considered significant at p<0.05 and *n* refers to the number of cells for electrophysiology experiments. All statistical analyses were conducted using GraphPad Prism 7.00 (GraphPad Software).

## RESULTS

### NMDA receptor activation evokes similar inward currents (I_NMDA_) in MNNs from sham and HF rats

We first evaluated whether the NMDAR-evoked current in MNNs differed between sham and HF rats. To this end, responses to varying durations of a focally applied NMDA (50 μM, 100–500 ms) were tested. As shown in Figure 1, I_NMDA_ increased in magnitude as a function of the puff duration both in MNNs from sham and HF rats (n=18 and 31, respectively, I_NMDA_ amplitude: p< 0.05 for both sham and HF respectively; I_NMDA_ area: p<0.05 for sham and HF respectively, RM one-way ANOVA). However, no differences were observed between sham and HF rats for the same NMDA puff duration (p>0.05). I_NMDA_ was almost completely eliminated in the presence of the NMDAR blocker AP5 (500 μM) (I_NMDA_ area 300 ms: Control 331.95 ± 42.97 nA*s vs AP5 0.062 ± 0.015 nA*s, p<0.0001, unpaired *t* test). We also did not observe any significant differences in the I_NMDA_ in identified GFP-VP MNNs from sham and HF rats (300 ms: sham 242.49 ± 62.68 nA*s vs HF 183.38 ± 89.69 nA*s, p>0.05, n =7 for both sham and HF, unpaired *t* test).

**Figure 1.**
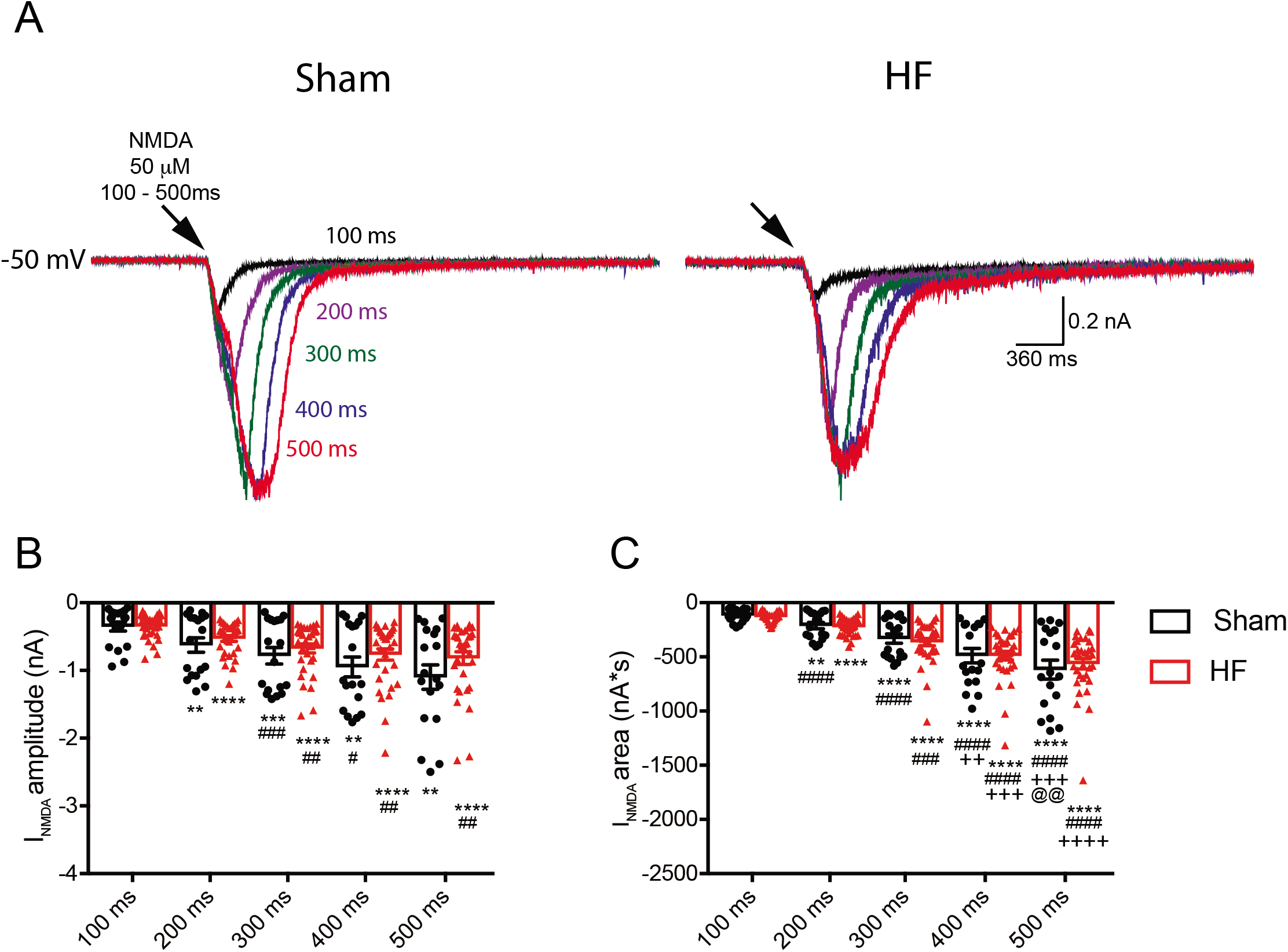
Focal application NMDA triggers an inward current (I_NMDA_) of similar magnitude in MNNs from Sham and HF rats. **A**, representative voltage-clamp recordings of MNNs from a Sham and a HF rat showing the responses of 50 μM NMDA focal application (100-500 ms) at −50mV. **B and C**, summary bar graphs showing the mean I_NMDA_ amplitude (B) and area (C) as a function of NMDA puff duration in MNNs from Sham (n=18) and HF (n=31) rats. **p<0.01, ***p<0.001 and ****p<0.0001 vs 100 ms. #p<0.05, ##p<0.01, ###p<0.001 and #### p<0.0001 vs 200 ms. ++p<0.01, +++p<0.001 and ++++p<0.0001 vs 300 ms. @@p<0.01 vs 400 ms. One-way ANOVA repeated measures.

### Blockade of SK channels increases I_NMDA_ magnitude in MNNs in sham but not HF rats

To asses a potential crosstalk between NMDARs and SK channels, we repeated similar experiments in the presence of the SK channel blocker apamin (200 nM). As shown in Figure 2, I_NMDA_ magnitude in MNNs of sham rats was significantly increased following application of apamin (I_NMDA_ amplitude: Control 0.504 ± 0.07 nA vs Apamin 0.658 ± 0.098 nA; I_NMDA_ area: Control 279.98 ± 33.64 nA*s vs Apamin 353.08 ± 41.39 nA*s, n=20, p<0.01 for both amplitude and area, RM two-way ANOVA). Importantly, the apamin-induced increase of I_NMDA_ in MNNs from sham rats was largely prevented in the presence of the Ca^2+^ chelator BAPTA (% change in I_NMDA_ – amplitude: 0.5 ± 7.8%; area: 4.9 ± 9.2%, p>0.05 for both amplitude and area, n=12, paired *t* test). These results suggest that the intracellular Ca^2+^ raise induced by NMDARs activation gates in a rapid and spatially localized manner SK channels, which in turn act in a negative feedback manner to dampen the evoked I_NMDA_.

**Figure 2.**
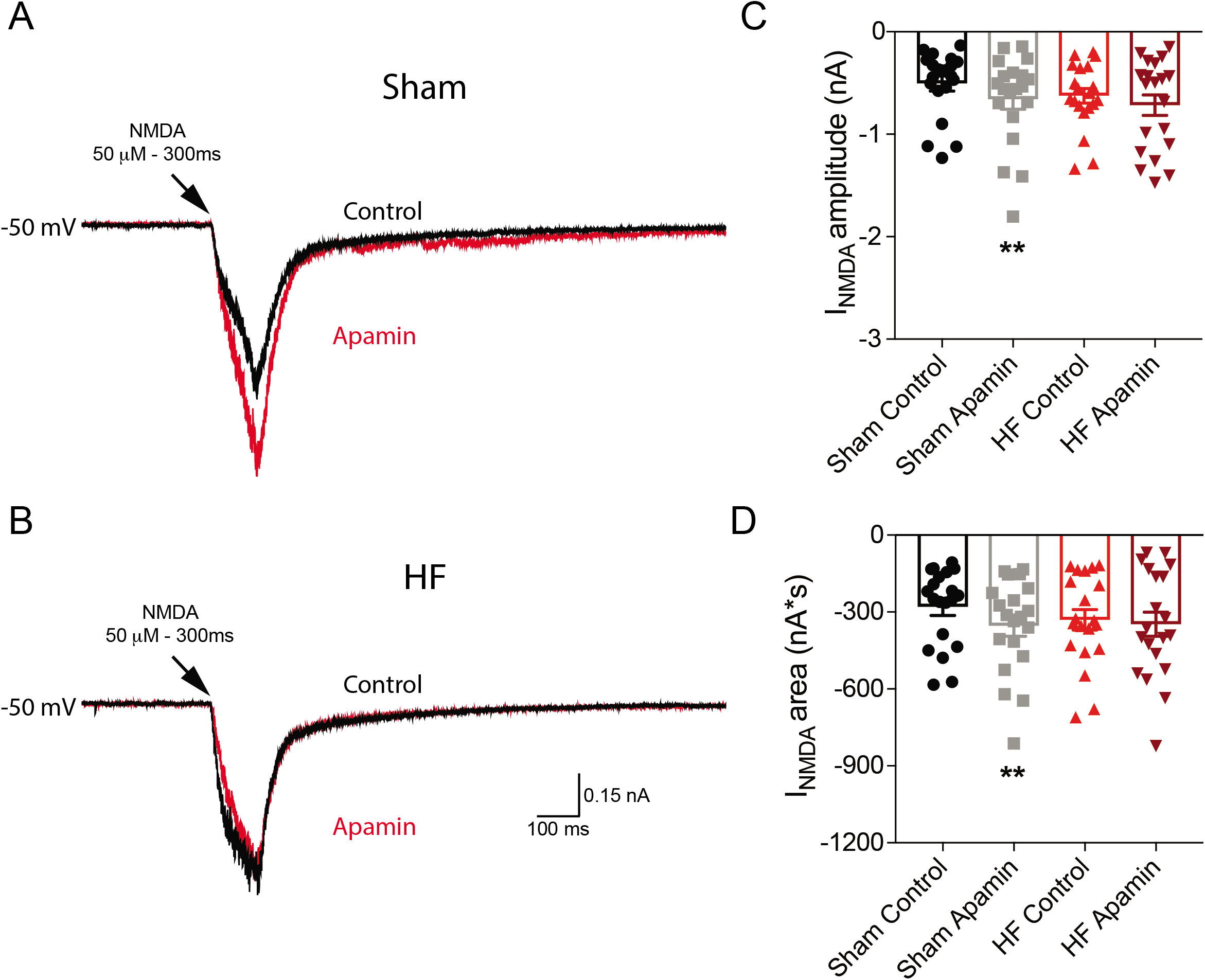
I_NMDA_ is increased by apamin in MNNs from Sham but not HF rats. **A and B**, representative voltage-clamp recordings of MNNs from a Sham (A) and HF (B) rat showing the responses of 50 μM NMDA focal application (300 ms) at −50 mV in the absence (control, black trace) and presence (red trace) of apamin (200 nM). **C and D**, summary bar graphs showing mean I_NMDA_ amplitude (C) and area (D) in MNNs from Sham (n=20) and HF (n=20) rats at −50 mV before and after apamin treatment. **p<0.001 vs Sham Control (paired *t* test).

Importantly, the effect of apamin was not observed in MNNs from HF rats (n=20, p>0.05 for both amplitude and area, RM two-way ANOVA, Fig.2), suggesting that the NMDA-SK channel coupling is blunted in MNNs of HF rats.

### NMDAR activation evokes a larger intracellular Ca^2+^ increase in MNNs in HF rats

To determine whether the blunted NMDAR-SK channel coupling in MNNs from HF rats was due to a diminished NMDAR-evoked increase in intracellular Ca^2+^ levels following NMDAR activation, we performed simultaneous voltage-clamp and fast confocal Ca^2+^ imaging in MNNs from sham and HF rats (n=17 and 30, respectively) that were loaded with the Ca^2+^ sensitive dye Fluo-5F. Representative recordings obtained from a MNNs in sham and HF rats are shown in Fig. 3. NMDAR activation (50 μM, 500 ms, Vm: −50 mV) evoked a large increase in intracellular Ca^2+^ that displayed slower rise and decay kinetics when compared to the underlying I_NMDA_. Thus, the peak Ca^2+^ increase occurred after the end of the NMDA puff, and the Ca^2+^ decay time course lasted several seconds beyond the end of the stimulation. Similar to the previous experiments, while the magnitude of I_NMDA_ did not differ between sham and HF rats (p>0.05 for both amplitude and area, unpaired *t* test, Fig. 3C and D), the magnitude of the NMDAR-evoked ΔCa^2+^ was significantly larger in MNNs from HF compared to sham rats (Fig. 3E and F; p<0.001 for both amplitude and area, unpaired *t* test). This difference persisted even when the magnitude of the evoked ΔCa^2+^ was normalized by the underlying I_NMDA_ (I_NMDA_ area: sham 0.016 ± 0.005 F/F_0_/nA*s vs HF 0.046 ± 0.011 F/F_0_/nA*s, p=0.05, unpaired *t* test). Taken together, these results suggest that the blunted NMDAR-SK channel coupling in MNNs from HF rats is not due to a diminished I_NMDA_-evoked ΔCa^2+^. In fact, a larger ΔCa^2+^ was observed in HF compared to sham rats.

**Figure 3.**
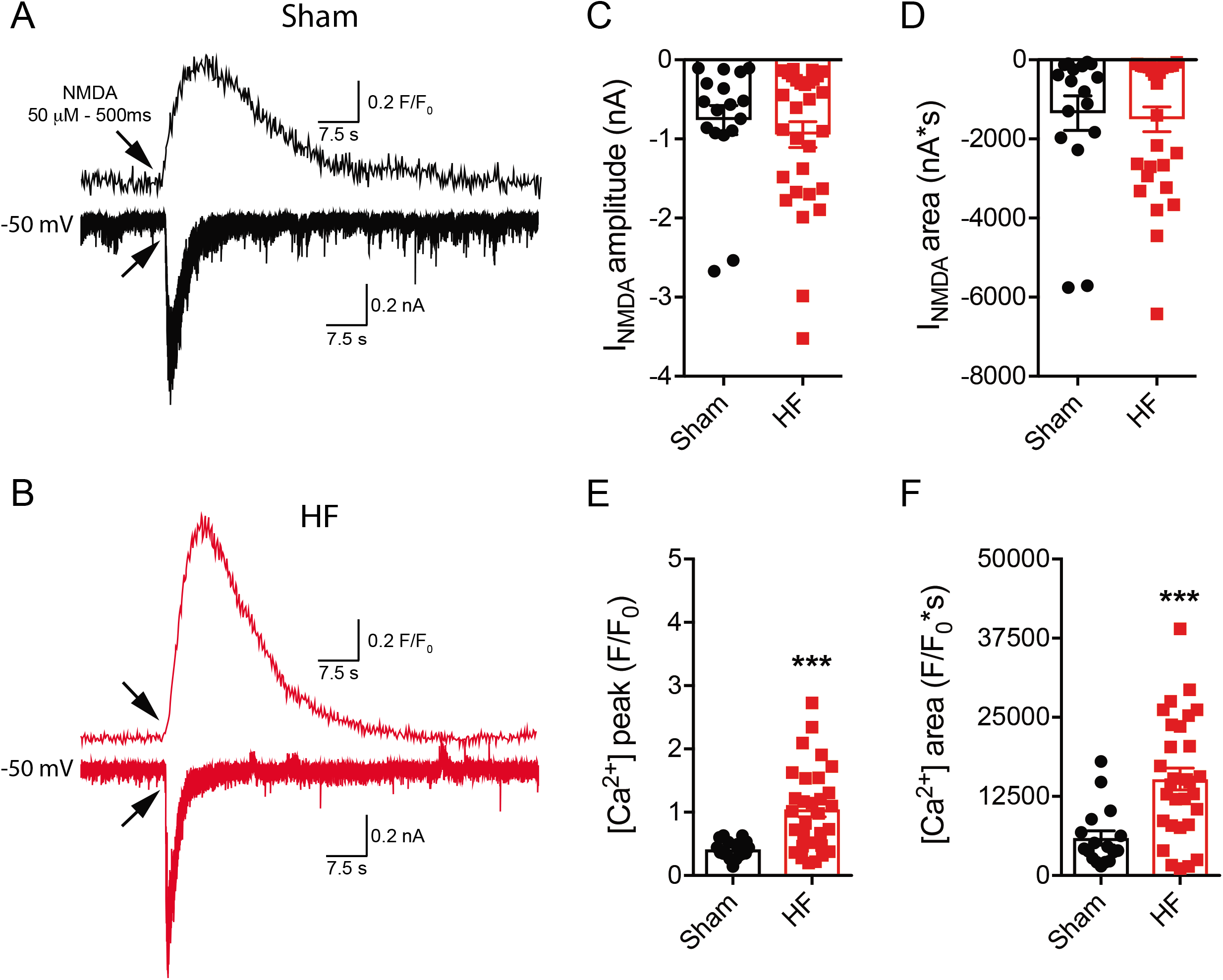
Representative samples of simultaneous recordings of I_NMDA_ and concomitant changes in intracellular Ca^2+^ levels in MNNs from Sham and HF rats. **A and B**, representative simultaneous voltage-clamp recordings of I_NMDA_ and Δ[Ca^2+^]_i_ in responses to a focal application of NMDA (50 μM, 500 ms) in MNNs from a Sham (A) and a HF (B) rat at −50 mV. **C and D**, summary bar graphs showing the mean amplitude (C) and area (D) of I_NMDA_ in MNNs from Sham (n=17) and HF (n= 30). **E and F**, summary bar graph showing mean Δ[Ca^2+^]_i_ peak (E) and area (F) induced by NMDA focal application in MNNs from Sham and HF rats. ***p<0.001 vs Sham (unpaired *t* test).

### Blockade of the SK channels potentiate NMDAR-evoked firing in MNNs in Sham but not HF rats

To investigate the functional relevance of the NMDAR-Ca^2+^-sensitive K^+^ channel crosstalk in MNNs, we performed current clamp experiments and assessed NMDAR evoked firing activity before and after blockade of SK channels. Moreover, we also compared whether SK channels differentially affected firing evoked by either direct current injection and NMDAR-evoked firing. Thus, each MNN was subjected both to a depolarizing current pulse (60 pA, 1 sec) and a puff of NMDA (50 μM, 300 ms), aiming to evoke ~15-20 spikes in each stimulation under basal conditions (i.e. before apamin). Representative examples for each stimulation modality evoked in MNNs from sham and HF rats are shown in Fig. 4.

**Figure 4.**
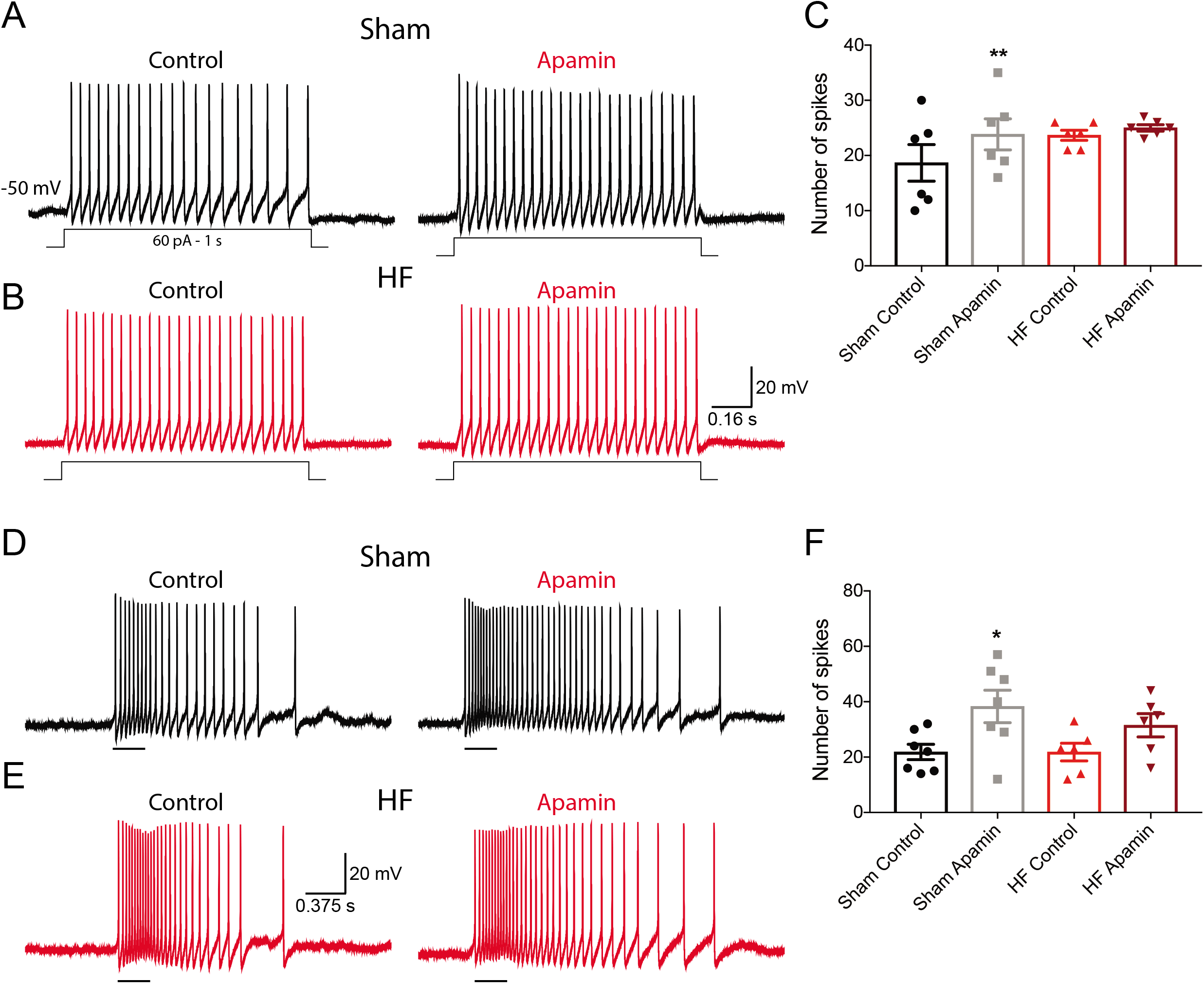
Effects of apamin on current injection- or NMDAR-evoked firing in MNNs from Sham and HF rats. **A and B**, representative current-clamp recordings from MNNs in Sham (A) and HF (B) rats showing responses to a depolarizing current injection (60 pA, 1 s) before and after apamin (200 nM). C, summary bar graph showing the mean number of action potentials evoked by depolarizing current injection before and after apamin treatment (n=6 and 6 in Sham and HF, respectively). **D and E**, representative current-clamp recordings from MNNs neurons in Sham (D) and HF (E) rats showing NMDA-evoked firing before and after apamin (200 nM) treatment (horizontal black bar under the traces: NMDA application – 50 μM, 300 ms). **F**, summary bar graph showing the mean number of action potentials evoked by NMDA focal application (50 μM, 300 ms) before and after apamin treatment (n=7 and 6 in Sham and HF, respectively). *p<0.05 and **p<0.01 vs Sham Control (paired *t* test).

In MNNs from sham rats, blockade of SK channels significantly increased the number of evoked spikes in response to both current injection (Fig. 4A and C, n=6, p<0.01, paired *t* test) and a puff of NMDA (Fig. 4D and F, n=6, p<0.05, paired *t* test). The degree of apamin effect was significantly larger in response to NMDAR-evoked firing (% change in firing: current injection 36.59 ± 12.55% vs NMDAR activation 93.69 ± 21.56%, p<0.05). Conversely, apamin failed to increase the number of evoked spikes in MNNs from HF rats, both in response to current injection (Fig. 4B and C, n=6, p>0.05, paired *t* test) as well as in response to a puff of NMDA, although a tendency for an effect was observed in the latter (Fig. 4E and F, n=6, p=0.06, paired *t* test). Importantly no differences in the input resistance of MNNs was observed between sham and HF rats (sham 605.28 ± 53.39 MΩ vs HF 501.36 ± 40.26 MΩ, p>0.05).

Taken together, these results support that a functional coupling between NMDARs and SK channels regulate membrane excitability and firing discharge in MNNs, and that this coupling is blunted in HF rats.

### The NMDAR-SK channel coupling regulates spike frequency adaptation during a burst of action potentials in MNNs from sham but not HF rats

To further investigate the impact of the NMDAR-SK channel coupling on the firing properties of MNNs, and given the importance of SK channels in regulating interspike intervals and spike frequency adaptation (SFA) (Alger & Nicoll, 1980; Hotson & Prince, 1980; Madison & Nicoll, 1984), we performed a detailed analysis of the effects of apamin on the interspike interval changes during direct current injection and NMDAR-evoked firing in both sham and HF rats (see *Methods*). In MNNs in sham rats, the interspike interval during a burst of action potentials evoked by direct current injection progressively increased as a function of the spike number in the train (Fig. 5A, p=0.01, RM one-way ANOVA, n=6), indicative of SFA. The degree of SFA, measured as the slope of a linear regression (Methods) was significantly diminished by apamin (p<0.05, paired *t* test, Fig. 5C).

**Figure 5.**
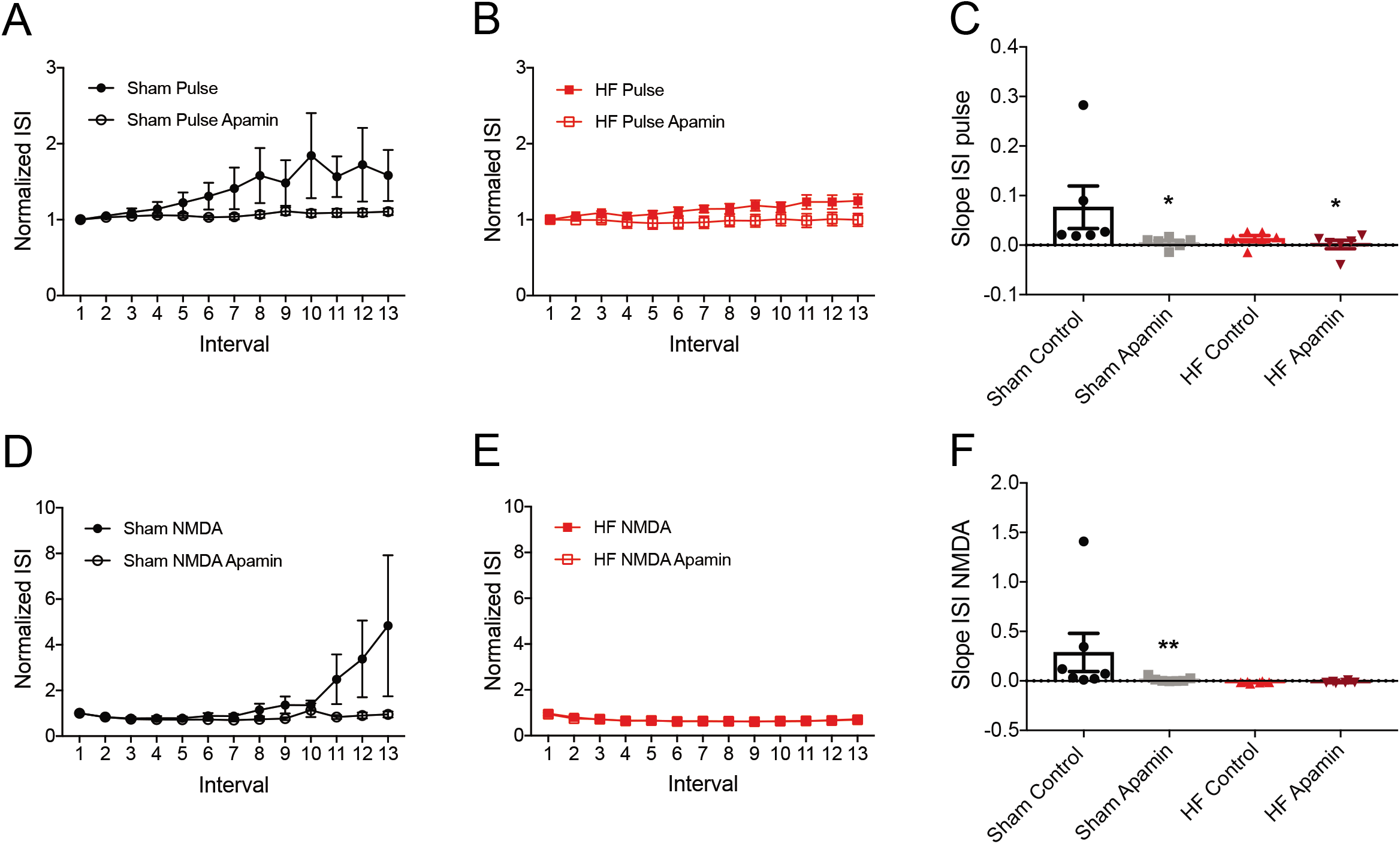
Effects of apamin on spike frequency adaptation following current injection-or NMDAR-evoked firing in MNNs from Sham and HF rats. **A and B**, mean plots of normalized ISI as a function of the number of the spike in the train following a depolarizing current injection (60 pA, 1 s) in MNNs from Sham (A) and HF (B) rats in the absence (control) and presence of apamin (200 nM). **C**, summary bar graph showing the mean slope of the ISI function in MNNs from Sham (n=6) and HF (n=6) rats in response to a depolarizing current injection. **D and E**, mean plots of normalized ISI as a function of the number of the spike in the train of NMDAR-evoked firing (50 μM, 300 ms) in MNNs from Sham (D) and HF (E) rats in the absence (control) and presence of apamin. F, summary bar graph showing the mean slope of the ISI function in MNNs from Sham (n=7) and HF (n=6) rats in response to NMDAR-evoked firing in MNNs from Sham and HF rats due to depolarizing current injection or NMDA puff. *p<0.05 and **p<0.01 vs Sham respective control (paired *t* test).

SFA during NMDAR-evoked firing was also present in MNNs in sham rats (Fig. 5D, p=0.05, RM one-way ANOVA) but displayed a clearly different pattern as that observed with direct current injection: An increase in the ISI duration was observed for later spikes in the train, and the degree of SFA was more pronounced, compared to that of observed in response to direct current injection. However, and similar to the latter, the degree NMDAR-evoked SFA was also significantly diminished in the presence of apamin (p<0.01, paired *t* test, Fig. 5F, n=7).

In MNNs of HF rats, SFA was still evident when spiking was evoked by direct current injection, as compared to sham rats (Fig. 5B, p<0.01, RM one-way ANOVA, although its degree was evidently less compared to sham rats (Fig. 5B). Apamin however still diminished the degree of SFA in HF rats (p<0.05, paired *t* test, Fig. 5C). Importantly, no SFA was observed in response to NMDAR-evoked firing in MMNs from HF rats (Fig. 5E). In fact, a significant progressive increase in ISI frequency was observed in this case (p<0.01, RM one-way ANOVA), as also supported by a negative slope in the relationship (Fig. 5F), which was not affected by apamin application (p=0.7, paired *t* test, Fig. 5F, n=6).

### The NMDAR-SK channel coupling influence membrane excitability in the absence of spiking activity

Experiments in voltage-clamp (Fig. 2) support that the coupling between NMDAR and SK channels can even occur in the absence of action potential firing. Thus, to determine whether this coupling was also sufficient to influence membrane potential depolarization evoked by NMDAR activation, we repeated a subset of current-clamp experiments in the presence of the voltage-gated Na^+^ channel blocker tetrodotoxin (TTX, 1 μM). As shown in Fig. 6, the magnitude and duration of the NMDAR-evoked membrane depolarization in MNNs from sham rats were significantly enlarged and prolonged, respectively, in the presence of apamin (n=15, p<0.01 and 0.05, respectively for depolarization area and duration, paired *t* test). Apamin, however, failed to affect the NMDAR-evoked membrane depolarization in MNNs from HF rats (n=12, p>0.05 for depolarization area and duration, paired *t* test).

**Figure 6.**
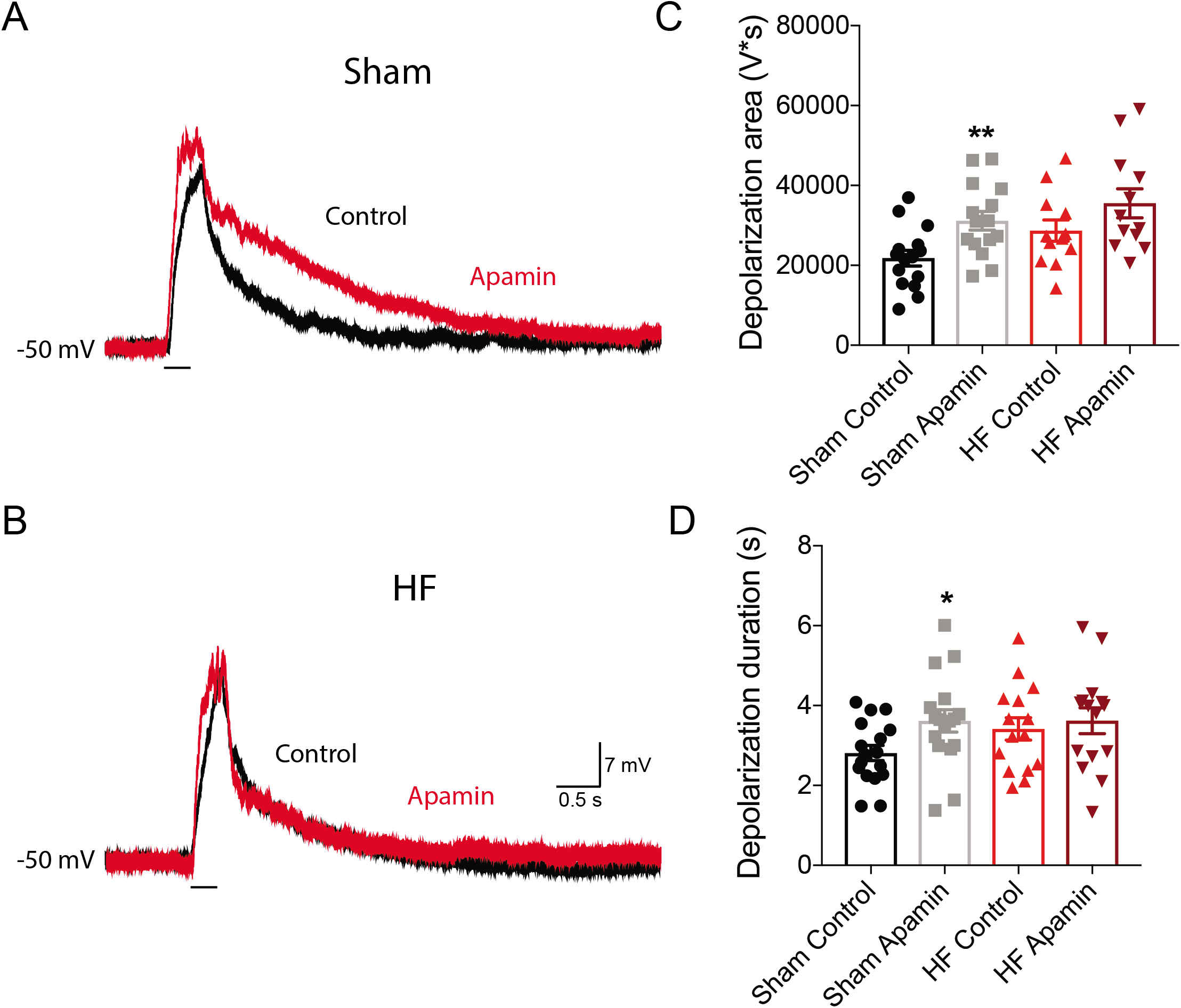
NMDA-evoked depolarization is augmented by apamin only in MNNs from Sham and rats. **A and B**, representative current-clamp recordings of MNN from a Sham (A) and a HF (B) rat showing the depolarization evoked by NMDA focal application (horizontal black bar under the traces: 50 μM, 300 ms) in the absence (control, black trace) and presence (red trace) of apamin (200 nM). **C and D**, summary bar graph showing the magnitude of NMDA-induced depolarization area (C) and duration (D) in MNNs from Sham and HF rats before and after apamin treatment (n=15 and 12 in Sham and HF, respectively). *p<0.05 and **p<0.01 vs Sham Control (paired *t* test or RM two-way ANOVA).

### Global intracellular Ca^2+^ chelation with BAPTA potentiates I_NMDA_ in MNNs of both sham and HF rat

Our results support a blunted NMDAR-SK channel coupling in MNNs from HF rats. Despite this, the basal I_NMDA_ magnitude in the latter was not significantly larger compared to those recorded in Sham rats. Based on this, along with our finding showing a larger NMDAR-evoked ΔCa^2+^ in HF rats, we hypothesize that a different Ca^2+^-sensitive mechanism could be activated to in MNNs of HF rats in order to compensate for the diminished contribution of SK channels. To test for this, we performed recordings in MNNs dialyzed with the Ca^2+^ chelator BAPTA. Given that cells were dialyzed with BAPTA from the beginning of the recording, we compared results to similar recordings obtained from separate cells using a normal internal solution. As shown in Fig. 7, chelation of intracellular Ca^2+^ with BAPTA significantly enhanced the magnitude of I_NMDA_ in MNNs from both sham (n=44 and 33 in control and BAPTA, respectively, p<0.05, unpaired t test) and HF rats (n=53 and 52 in Control and BAPTA, respectively, p<0.01 and p<0.05 for amplitude and area, respectively unpaired t test). Importantly, the degree of I_NMDA_ enhancement evoked by apamin or BAPTA in MNNs from sham rats was not significantly different (28 ± 5 and 42 ± 18% increase in I_NMDA_ area, respectively, p>0.05). Moreover, as reported above, dialyzing MNNs in sham rats with BAPTA occluded the apamin-evoked potentiation of I_NMDA_. Taken together, these results suggest that SK channels constitute the main Ca^2+^-sensitive mechanism regulating the magnitude of I_NMDA_ in sham rats, and that in rats with HF, a non-SK, Ca^2+^-sensitive mechanism may act as a compensatory mechanism to the blunted action of SK channels in this experimental group.

**Figure 7.**
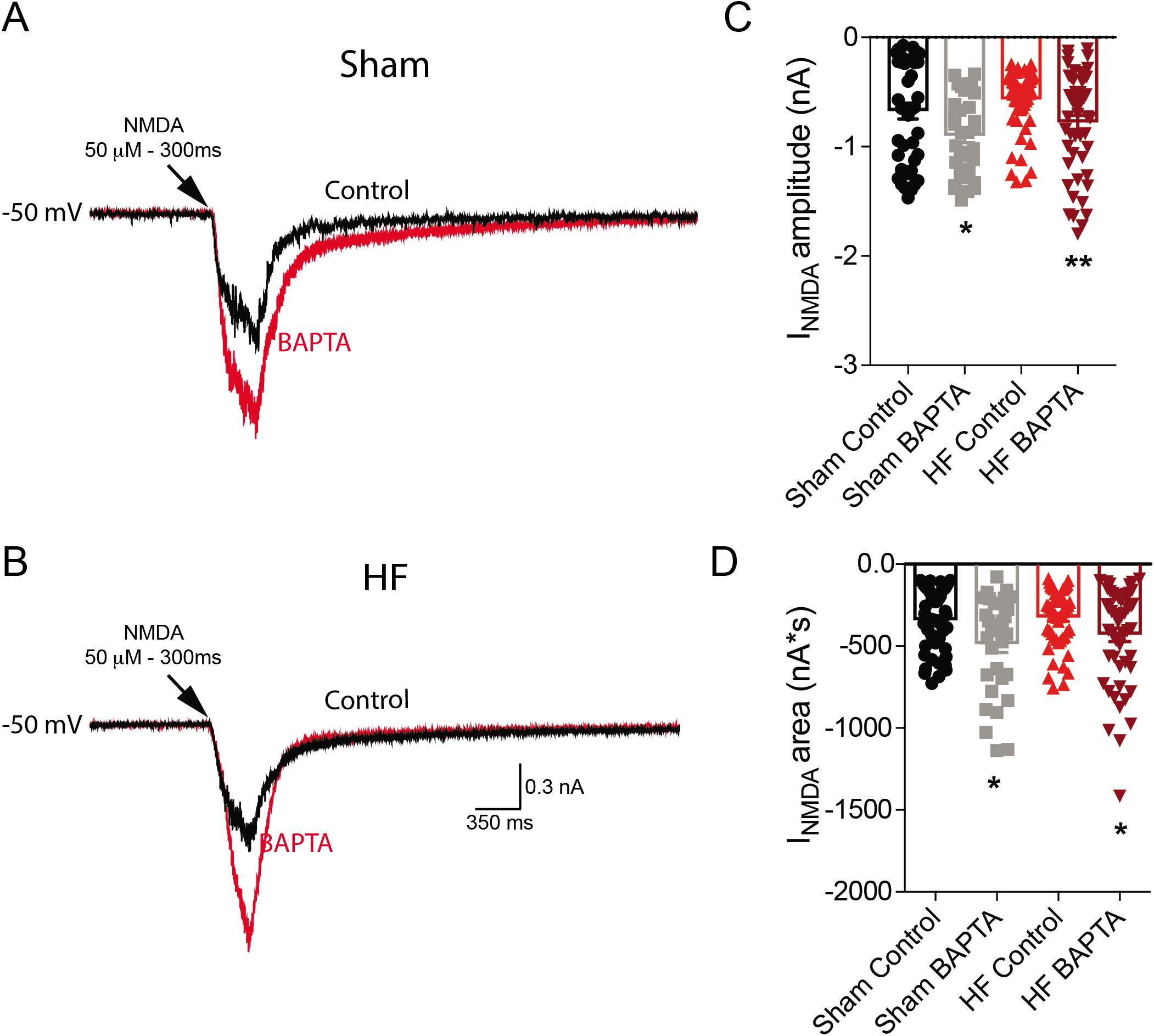
Ca^2+^ chelation promotes an enhancement of I_NMDA_ in MNNs from Sham and HF rats. **A and B**, representative voltage-clamp recordings of MNN from a Sham (A) and HF (B) rat in control condition (black trace) and with BAPTA (10 mM, red trace) added to the internal solution in the recording pipette showing the responses of NMDA focal application (50 μM, 300 ms) at −50 mV. **C and D**, summary bar graphs showing the changes induced by BAPTA in I_NMDA_ amplitude (C) and area (D) in MNNs from Sham (n=44 and 33 in control and BAPTA, respectively) and HF rats (n=53 and 52 in control and BAPTA, respectively). *p<0.05 and **p<0.01 vs respective control (unpaired *t* test).

## DISCUSSION

Small conductance Ca^2+^-sensitive K^+^ channels (SK) are ubiquitously expressed in a variety of CNS neurons, including MNNs (Greffrath *et al*., 2004). SK channels are typically gated by an increase in intracellular Ca^2+^ levels following opening of voltage-gated Ca^2+^ channels (VGCC) during a burst of action potentials (Bond *et al*., 1999). In addition to VGCC, glutamate NMDARs constitute another major source of Ca^2+^ in CNS neurons. In fact, recent studies in hippocampal and cortical neurons showed that SK channels can be gated in a Ca^2+^-dependent manner following NMDAR activation (Ngo-Anh *et al*., 2005; Babiec *et al*., 2017), and that this coupling plays an important role in regulating synaptic efficacy and plasticity. Still whether a direct interaction between NMDARs and SK channels also occurs in MNNs, and if so, what the functional relevance of this coupling is, remains until now unknown.

### Negative feedback loop between NMDARs and SK channels regulate membrane excitability and firing discharge in MNNs

One of the major findings from this study is that under control conditions, NMDARs in MNNs are functionally coupled to SK channels, forming a negative feedback loop that restrains the excitatory effect on membrane potential and firing activity that follows activation of NMDARs. This conclusion is supported by several lines of evidence that emerged from our studies. Firstly, we found that the magnitude of an inward excitatory current evoked by NMDAR activation (I_NMDA_) was significantly enhanced following the specific inhibition of SK channels with apamin. The fact that the I_NMDA_ in the presence of apamin was potentiated promptly during the time course of the short NMDA application supports a very fast, and spatially localized interaction between NMDARs and SK channels. This is also supported by the fact that intracellular dialysis with the high affinity Ca^2+^ chelator BAPTA prevented the effect of apamin on I_NMDA_, as also described in Ngo-Anh and colleagues (Ngo-Anh *et al*., 2005), supporting a putative Ca^2+^ microdomain linking NMDARs and SK channels. Importantly, BAPTA *per se* increased the magnitude of I_NMDA_, to a similar extent as that evoked by apamin. Given that MNNs express other Ca^2+^-activated K^+^ sensitive channels, including BK channels (Greffrath *et al*., 2004; Ohbuchi *et al*., 2010), these results support that at least under control conditions, SK channels constitute the main type of channel involved in this rapid Ca^2+^-sensitive inhibitory feedback loop triggered by activation of NMDARs.

While we did not attempt to identify the phenotype of each recorded MNN, we found no evidence of two populations differentially affected. Moreover, a subset of experiments performed in eGFP-VP rats (Ueta *et al*., 2005) indicated that the effects were observed in identified VP neurons. Thus, while our results suggest that the NMDAR-SK channel coupling is present in both VP and OT neurons, we could conclusively say that it is present in VP neurons.

To determine whether the NMDAR-SK channel inhibitory feedback loop was functionally relevant, we performed current clamp experiments. Our results show that the total number of action potentials evoked by NMDAR activation was significantly enhanced following SK channel blockade, supporting that the NMDAR-SK channel coupling acts in a rapid manner to efficiently restrain the NMDAR-mediated increase in firing discharge. Previous studies have shown that in MNNs activation of SK channels during repetitive action potential firing contributes to spike frequency adaptation (SFA), i.e., a progressive increase in the interspike interval duration (Greffrath *et al*., 2004). We report here that NMDAR-evoked firing in MNNs also results in SFA, which was almost completely eliminated by apamin, being thus another line of evidence supporting the functional coupling between NMDARs and SK channels in MNNs.

Importantly, and as previously described (Ferreira-Neto *et al*., 2017), we found that apamin also increased the number of action potentials evoked by direct current injection. However, for the same number of action potentials evoked, the effect of apamin was markedly greater when action potentials were evoked by NMDAR activation (i.e., ~36% vs 94% increase when comparing direct current injection with NMDAR activation, respectively). These results support that compared to VGCC, NMDARs couple more efficiently to SK channels. This is likely the result of differences in the spatiotemporal properties of the Ca^2+^ dynamics evoked by NMDARs and VGCC, differences that may include a larger amount of Ca^2+^ entry per spike, or closer spatial relationship of NMDARs to SK channels. Future experiments will be needed to further elucidate this phenomenon.

In a series of recent publications, we showed that functional extrasynaptic NMDARs (eNMDARs) play an important role in mediating the excitatory effect of glutamate in MNNs (Fleming *et al*., 2011; Potapenko *et al*., 2012, 2013; Stern & Potapenko, 2013; Zhang *et al*., 2017). Moreover, and as previously showed in other brain regions (Hardingham & Bading, 2010), eNMDARs in MNNs are differentially coupled to distinct intracellular signaling cascades and targets. Thus, we found eNMDARs (but not synaptic ones) to be negatively and positively coupled, respectively, to A-type K^+^ channels (Naskar & Stern, 2014; Zhang *et al*., 2017) and GABA_A_ receptors (Potapenko *et al*., 2013). Thus, it will be important to determine in future studies the relative contribution of synaptic and eNMDARs to the functional coupling to SK channels described herein.

### Blunted NMDAR-SK channel coupling in MNNs from HF rats

Another important finding from these studies is that the negative feedback loop between NMDARs and SK channels was largely blunted or absent in MNNs from HF rats. Thus, we found that in this experimental group apamin failed to increase I_NMDA_, to enhance the NMDAR-evoked membrane depolarization and the number of evoked action potentials as observed in MNNs from sham rats.

Several mechanisms could contribute to the blunted NMDAR-SK channel interaction in HF, including a diminished I_NMDA_. This however, was not the case. In fact, given the blunted inhibitory feedback, a larger I_NMDA_ would be expected in MNNs from HF rats (see more below about this). A diminished NMDAR-ΔCa^2+^ could also lead to diminished activation of SK channels. However, and as we previously reported (Stern & Potapenko, 2013) we found a larger NMDAR-ΔCa^2+^ in MNNs from HF. Finally, changes in SK channel function/expression could also contribute to the blunted negative feedback reported in this study. In this sense, we recently reported a diminished SK2/SK3 channel subunit mRNA expression in the SON of HF rats (Ferreira-Neto *et al*., 2017), suggesting that a reduction in the number of available SK channels could constitute a key underlying mechanism. However, whether the global decrease in SK channel mRNA expression within the SON specifically affected the availability of SK channels in the vicinity of NMDARs is at present unknown. Moreover, we cannot rule out at present whether a diminished SK channel Ca^2+^-sensitivity also contributes to the altered NMDAR-SK channel crosstalk during HF. Thus, future studies are warranted to explore more specifically the relative contribution of these potential mechanisms.

As stated above, and based on a blunted NMDAR-SK negative feedback in HF, we would have expected a larger baseline I_NMDA_ in MNNs of HF rats. However, while a tendency was observed, this difference was not statistically significant. A possible explanation for this apparent discrepancy could be explained by the results obtained from the BAPTA experiments. Thus, while SK channel blockade with apamin enhanced I_NMDA_ only in MNNs from sham rats, we found that global Ca^2+^ chelation with BAPTA potentiated I_NMDA_ in both experimental groups. Thus, one likely interpretation for these results is that a non-SK, Ca^2+^-sensitive mechanism is specifically activated in MNNs from HF rats, acting as a compensatory mechanism to the blunted action of SK channels in this condition. The molecular identity of this Ca^2+^-sensitive mechanism, and the time course relationship between its onset and the onset/evolution of the disease remains to be established.

Taken together, our studies support a novel functional crosstalk between NMDARs and SK channels in MNNs that acts as an important negative feedback mechanism that regulates neuronal excitability and activity in the SON. Moreover, our studies support that a blunted crosstalk may contribute to the pathophysiology of neurohumoral activation during HF.

## ADDITIONAL INFORMATION

### Authors contributions

Ferreira-Neto HC: performed surgeries, echocardiogram and experiments, analyzed data, interpreted results of experiments, prepared figures, drafted manuscript and approved final version of manuscript.

Stern IE: conception and design of the experiments, interpretation of data, edited, revised and approved final version of manuscript.

All authors approved the final version of the manuscript.

## ABBREVIATIONS

HF: heart failure;
IR-DIC: infrared differential interference contrast;
ISI: interspike interval;
MNNs: magnocellular neurosecretory cells;
NMDAR: NMDA receptors;
OT: oxytocin;
PVN: paraventricular nucleus;
SFA: spike frequency adaptation;
SK: small conductance Ca^2+^-activated K^+^ channels;
SON: supraoptic nucleus;
VP: vasopressin

